# NRPS genes provide antifungal activity to new *Bacillus* and *Paenibacillus* strains and change their expression in the presence of phytopathogens in a strain-specific manner

**DOI:** 10.1101/2023.12.06.570468

**Authors:** N.G. Vasilchenko, M.P. Kulikov, E.V. Prazdnova, A.V. Gorovtsov, A.V. Usatov, V.A. Chistyakov

## Abstract

Fusarium diseases pose a significant threat to agricultural crops. In agricultural practice, various chemical fungicides, typically triazole-based, are often used to control fusariosis. However, this practice contributes to the development of resistant strains among the pathogenic fungi. A potential alternative to chemical fungicide treatments involves utilizing biopreparations derived from natural antagonists of *Fusarium*. *Bacillus* and *Paenibacillus* bacteria, isolated from farmland soils have shown notable antagonistic activity against phytopathogenic *Fusarium* fungi. Genetic analysis via PCR has indicated the presence of nonribosomal peptide synthase genes in the isolated bacterial strains, with the absence of chitinase activity suggesting that the antifungal effect primarily stems from nonribosomal synthesized peptides. Furthermore, the expression of these genes is heightened in the presence of fungi during co-culturing, although the magnitude of this effect appears to be specific to each strain and dependent on the pathogen in question. Field trials have demonstrated the efficacy of a biocontrol preparation made from studied *Bacillus* and *Paenibacillus* strains. These results suggest promising prospects for the development of pathogen-targeted treatments.

## 1. Introduction

The vast majority of *Fusarium* species are plant pathogens capable of causing significant damage to agricultural crops, causing significant resulting in up to 40% losses in commodity grains and a decrease in overall crop quality (Ekwomadu et al., 2021; Nikitin et al., 2023).

Various chemical fungicides are used to control fusariosis in agricultural practice. The most common are triazole-based preparations, as well as combinations of triazoles with other active substances - prochloraz or spiroxamine (Chen et al., 2020; Feksa et al., 2019; He et al., 2023; Li et al., 2022). The prevalent use of triazoles in crop production poses a serious risk, as prolonged exposure to substances from the same class can lead to the development of resistant forms among pathogens (Yerkovich et al., 2020). This phenomenon is also typical for agricultural use of fungicides. The emergence of powdery mildew resistant to triazoles and several other fungicides was first noted back in 1987 (Weete and Wise, 1987), and resistant forms are currently observed in a number of other plant diseases (Resendiz Sharpe et al., 2018).

Bacteria from the *Bacillus* and *Paenibacillus* genera exhibit various antifungal mechanisms, which makes them promising candidates for the development of biopreparations for the control of fungal phytopathogens (Soni and Keharia, 2021). This group of aerobic spore- forming bacteria demonstrate strong antagonistic activity against various organisms: gram- negative and gram-positive bacteria (Olishevska et al., 2019; Soni and Keharia, 2021), a wide range of fungal species (Costa et al., 2022; Olishevska et al., 2019; Vasilchenko et al., 2022) and some insects (Ruiu, 2020). Such a wide range of antagonistic activities is determined by the synthesis of various enzymes and small functional molecules. Enzymes such as chitinases and beta-glucanases play a role in their antifungal properties by breaking down cell walls (Swiontek Brzezinska et al., 2020; Thakur et al., 2022). These bacteria also produce a variety of low molecular weight compounds, including nonribosomal peptides (NRPs), polyketides (PKs), and volatile organic compounds (VOCs) (Li et al., 2020; Olishevska et al., 2019).

NRPs are of particular interest for both fundamental and applied research, as new promising antibiotics, antimycotics and biosurfactants are constantly being discovered in this group of compounds (Olmedo et al., 2022; Ranjan et al., 2023). These molecules are produced via activity of nonribosomal peptide synthetases (NRPS), therefore, we can estimate the ability of bacteria to produce NRPs by studying NRPS genes.

This study aimed to investigate the role of NRPS genes and their expression in the antifungal effect of the 10 newly identified strains of *Bacillus* and *Paenibacillus* genera, and to test the prototype of antifungal preparation based on these strains. It was found that the expression of these genes increases when co-cultured with *Fusarium* strains in a strain-specific manner, which indirectly confirms their involvement in this activity.

## 2. Methods

### 2.1. Strains

Cultures of fungal pathogens *Fusarium oxysporum* (strains Ras6.1 and Ras6.2.1) and *Fusarium graminearum* (strains Fus1 and Fus5), strains with strong antifungal activity *Paenibacillus polymyxa* (K1.14, O1.27, O2.11, R3.13, R4.5, R4.24, R5.31, R6.14), *Bacillus velezensis* strains (V3.14, R4.6) and strains with weak activity (listed below in the order from 1 to 20) were isolated from soils of agricultural fields of Krasnodar region, Russian Federation. Strains with low activity were not used in the preparation for field experiments, but were used in laboratory experiments as a negative control group.

### 2.2. Primary screening of antagonistic strains

For initial selection of antagonist strains of *Fusarium*, we used the agar plate method of co-cultivation on nutrient medium (0.5 Malt extract agar: 0.5 nutrient agar), pre-inoculated with culture of either *F. oxysporum* or *F. graminearum* by plating the inoculum on the agar medium, then a pasteurized soil suspension was plated on top of the fungal culture on the same Petri dish. The double culture dishes were incubated for 3-5 days at 27-28°C. All bacteria around which a distinct zone of fungal growth suppression was formed were isolated and tested against *F. graminearum* (Fus1 and Fus5) and *F. oxysporum* (Ras6.1, Ras6.2.1). Only strains capable of suppressing all 4 strains of *Fusarium* were selected.

### 2.3. Determination of chitinase activity of antagonistic strains

To determine the chitinase activity of the strains, the strains were inoculated on chitin agar according to the method of Rishad (2017). The strains were cultured for 3 days at 27 °C. The presence or absence of this activity was recorded by the presence of a zone of clarification of the nutrient medium around the colony of the microorganism.

### 2.4. Identification of bacterial and fungal strains

The isolated phytopathogenic fungi and ten selected most active strains of bacteria- antagonists of phytopathogenic fungi were subjected to identification based on DNA analysis in the Russian National Collection of Industrial Microorganisms.

Identification was based on sequencing of 16s rRNA gene sequences and further phylogenetic analysis in the case of bacterial strains. Identification of phytopathogenic fungi was based on sequencing and phylogenetic analysis of 18s rRNA, 5.8s rRNA and internal transcribed spacer genes *ITS1* and *ITS2*. Detailed results are described in our previous paper (Gorovtsov et al., 2019).

### 2.5. Extraction of nucleic acids and cDNA production in this study

DNA extraction from overnight bacterial cells culture grown on Malt extract-Meat peptone agar was performed using "DNA Sorb С-М" extraction kit (Amplisens, Russia). The DNA was purified using "CleanMag DNA kit" (Eurogen, Russia).

RNA isolation from bacterial cells was performed using a "Extract RNA" kit (Eurogen, Russia) and them purified with "CleanRNA Standard" kit (Eurogen, Russia). The isolation was performed from bacterial culture grown on Malt extract- nutrient agar in co-culture with fungi and without co-culture as a control.

The production of complementary single-stranded DNA on RNA matrix was performed using the "MMLV RT kit" (Eurogen, Russia). The concentration and quality of isolated DNA, RNA and synthesized cDNA were measured using a spectrophotometer Nano Photometer P330 (Implen, USA) and QuantiFluor ST Fluorometer (Promega, USA).

### 2.6. NRPS gene expression analysis

The presence of NRPS genes was determined by PCR using a commercial kit "ScreenMix" (Eurogen, Russia) and DNA as a matrix. PCR results were visualized in 1.5% agarose gel with ethidium bromide and Gel-Doc gel-documentation system (Bio-Rad, USA). The primers presented in Table 1 were used to search for NRPS genes. Optimal temperatures for primer annealing were estimated by a gradient experiments.

**Table 1.**
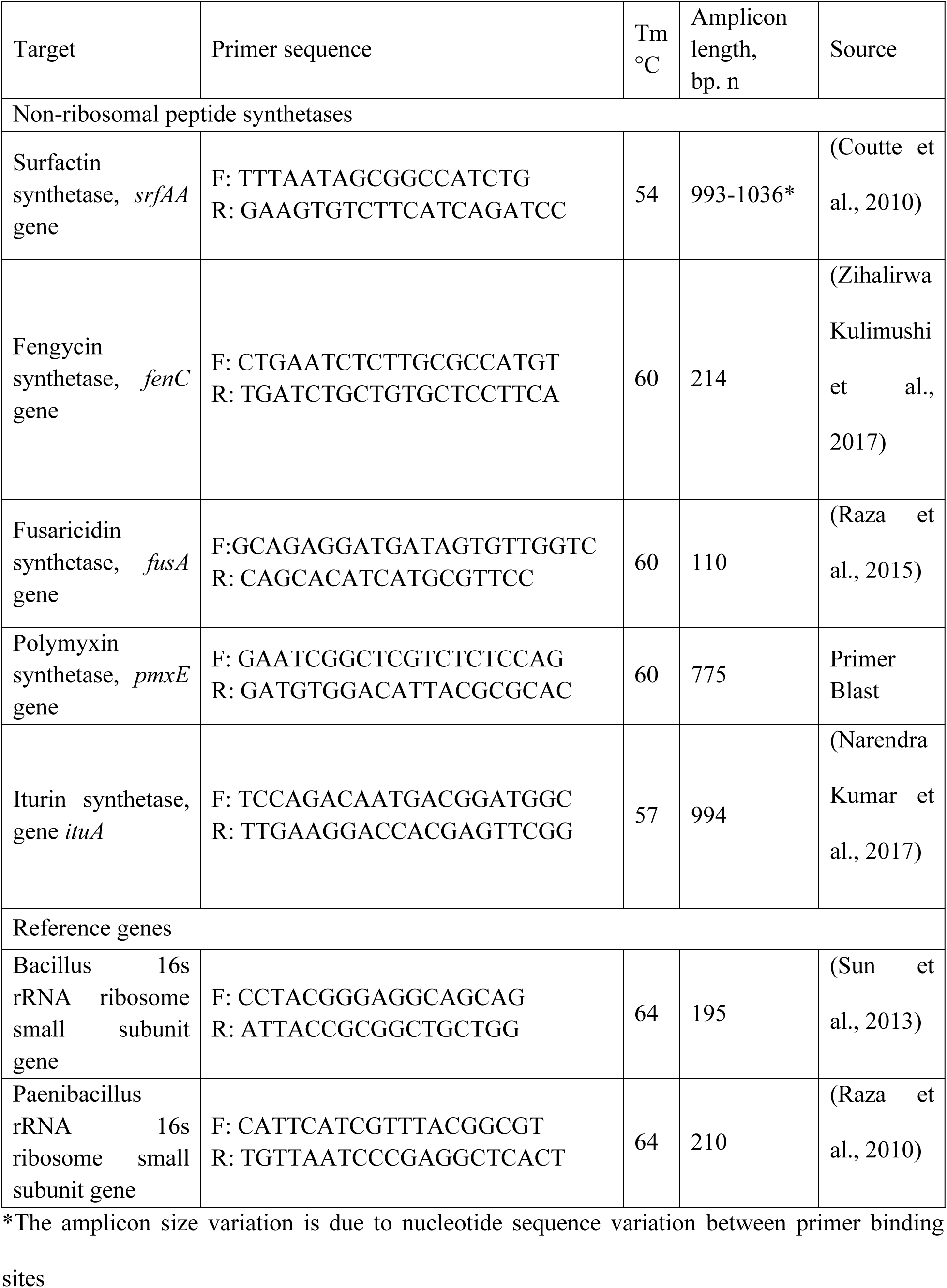
Characteristics of primers used to search for NRPS genes.

The expression levels of NRPS genes were assessed using a CFX96 Touch (Bio-Rad, USA) and a “PCR-mix SYBR Green I kit” (Syntol, Russia).

To study the effect of fungi presence on the change of NRPS gene expression patterns, fungi and bacteria were co-cultured on solid nutrient medium. A full description of the isolation and processing conditions is presented in section 2.5.

The changes in relative gene expression in antagonist bacterial strains in the presence of different *Fusarium* species were studied separately. Experiments were performed in 3 biological replicates.

The ***2^-ΔΔCT^*** method (Rao et al., 2013) was used to assess the relative expression level. 16s rRNA was used as a reference gene. The obtained values of relative gene expression were presented as arithmetic mean and standard error, which were further used for statistical calculations in R using ANOVA with post-hoc Tukey HSD test.

### 2.7. Testing the antifungal efficacy of a preparation in field studies

In a series of laboratory experiments, a biopreparation based on culture fluid containing 10 selected strains described above and their secondary metabolites actively inhibiting the growth of *Fusarium* was obtained.

Tests of the preparation were conducted on test fields 1-4 located in the territory of Krasnodar region, Russian Federation. The soil of the test fields is classified as chernozemic. Experiments with the application of the biopreparation were conducted according to the following scheme. Beet leaves were sprayed once and twice with biopreparation in liquid form. The application rate was 20 ml/ha. The number of viable bacteria in the preparation was 1·10^10^ CFU/mL. Single treatment of plants was carried out at the fork stage. The second treatment (in case of double treatment) was performed at the stage of leaf closure in the inter-rows.

Plant sampling was done with 5 samples from each field. To assess the infestation, root crops were and treated with 70% ethanol. A sterile knife was used to cut 3 fragments of 0.5 cm ^3^ from the center of each root crop, after which the fragments were placed on the surface of Chapek’s nutrient medium under sterile conditions. At the plant sampling sites, soil was removed from the 0-10 cm layer for further evaluation of the infection load level. To assess the level of infection load in soils, surface inoculation was performed from serial dilutions. Counting was carried out on the 5-7th day according to the morphology of colonies with confirmation of belonging to the genus *Fusarium* using microscopy.

Statistical analysis was carried out using the nonparametric Mann-Whitney U-criterion for pairwise comparisons to compare data pertaining to the sugar beet plant infestation and presence of *Fusarium* in the soil.

## 3. Results

### 3.1. Characterization of selected strains

Based on the results of the screening for antifungal activity, 10 bacterial strains were selected: 2 strains of the genus *Bacillus* (V3.14, R4.6) and 8 strains identified as bacteria of the genus *Paenibacillus* (K1.14, O1.27, O2.11, R3.13, R4.5, R4.24, R5.31, R6.14). The strains are described in more details in our previous paper (Gorovtsov et al., 2019).

### 3.2. Chitinase activity detection

In order to identify one of the prevalent mechanisms of antifungal activity, namely chitinase activity (Chang et al., 2010), a test was carried out on antagonistic bacterial strains. It was observed that none of the 10 selected bacterial strains exhibited chitinase activity, as indicated by the absence of a hydrolysis zone on colloidal chitin-containing nutrient medium during surface growth.

### 3.3. NRPS gene screening

An examination of NRPS genes was conducted. The results of genes search in the selected antagonistic bacterial strains are presented in Table 2. As can be seen from these data, *Bacillus* strains (V3.14, R4.6) contain the *fusA* gene, responsible for fusaricidin synthetase synthesis, alongside genes for surfactin (*srfAA*) and fengycin synthesis (*fenC*). While the presence of fusaricidin synthetase genes typically associated with *Paenibacillus* bacteria, its occurrence in *Bacillus* could be attributed to horizontal gene transfer or the presence of ortholog of this gene. The reference gene for the 16s rRNA subunit (*16sBam*), used as a positive control, was consistently detected in all strains.

**Table 2.** Genes of different non-ribosomal peptide synthetases detected in bacterial strains by PCR and electrophoresis methods.

|  |  |  | Gene |  |  |  |  |  |  |
| --- | --- | --- | --- | --- | --- | --- | --- | --- | --- |
|  |  |  | *srfAA* | *fenC* | *fusA* | *pmxE* | *ituA* | *16s Bam* | *16s Pmx* |
| Bacterial strains | (symbols in the graphs below) | 1 | - | - | - | - | - | + | - |
|  |  | 2 | - | - | - | - | - | + | - |
|  |  | 3 | - | - | + | - | - | + | - |
|  |  | 4 | - | - | - | - | - | + | - |
|  |  | 5 | - | + | - | - | - | + | - |
|  |  | 6 | - | - | - | - | - | + | - |
|  |  | 7 | - | - | - | - | - | + | - |
|  |  | 8 | - | - | - | - | - | + | - |
|  |  | 9 | + | - | - | - | - | + | - |
|  |  | 10 | - | - | - | - | - | + | - |
|  |  | 11 | - | - | - | - | - | + | - |
|  |  | 12 | - | - | - | - | - | + | - |
|  |  | 13 | - | - | - | - | - | + | - |
|  |  | 14 | - | - | - | - | - | + | - |
|  |  | 15 | - | - | - | - | - | + | - |
|  |  | 16 | - | - | - | - | - | + | - |
|  |  | 17 | - | - | - | - | - | + | - |
|  |  | 18 | - | - | - | - | - | + | - |
|  |  | 19 | - | - | - | - | - | + | - |
|  |  | 20 | - | - | - | - | - | + | - |
|  |  | K1.14(VI) | - | - | + | + | - | + | + |
|  |  | O1.27(I) | - | - | + | + | - | + | + |
|  |  | O2.11 (VIII) | - | - | + | + | - | + | + |
|  |  | R3.13 (II) | - | - | + | + | - | + | + |
|  |  | V3.14 (IX) | + | + | + | - | - | + | - |
|  |  | R4.5 (V) | - | - | + | + | - | + | + |
|  |  | R4.6 (IV) | + | + | + | - | - | + | - |
|  |  | R4.24 (X) | - | - | + | + | - | + | + |
|  |  | R5.31 (III) | - | - | + | + | - | + | + |
|  |  | R6.14 (VII) | - | - | + | + | - | + | + |

\* "+" - presence of the gene, "-" - absence of the gene

Within *Paenibacillus* strains, only genes of fusaricidin and polymyxin (*pmxE*) synthetases were detected among the studied targets. These genes were present in all 8 *Paenibacillus* strains. Notably, strains 1 - 20 exhibited minimal antagonistic activity against the fungus *F. graminearum* Fus1. As can be seen from the table 2, only 3 weak antagonistic strains (number 3,5 and 9) were found to possess NRPS genes. Each of these strains had only a single NRPS gene detected.

### 3.4. Analysis of NRPS gene expression in bacterial antagonistic strains and strains with weak antagonistic activity against *Fusarium*

The analysis of NRPS genes expression was conducted in bacterial strains exhibiting high antagonistic activity against *Fusarium*, as well as strains with low activity but containing NRPS genes. For this experiment, bacterial were cultured both in the presence and absence of *F. oxysporum* Ras6.2.1.

While some weakly antagonistic bacteria shared the same genes as their highly antagonistic counterparts, they did not exhibit activation of these genes in the presence of the fungus *F. oxysporum* Ras6.2.1. Conversely, significant activation of NRPS gene expression in the presence of fungus was observed in highly antagonistic bacterial strains. Thus, it was determined that activation of NRPS gene expression in the presence of fungi is specific to bacterial strains with high antagonistic activity. The summarized results are presented in Figure 1.

**Fig. 1.**
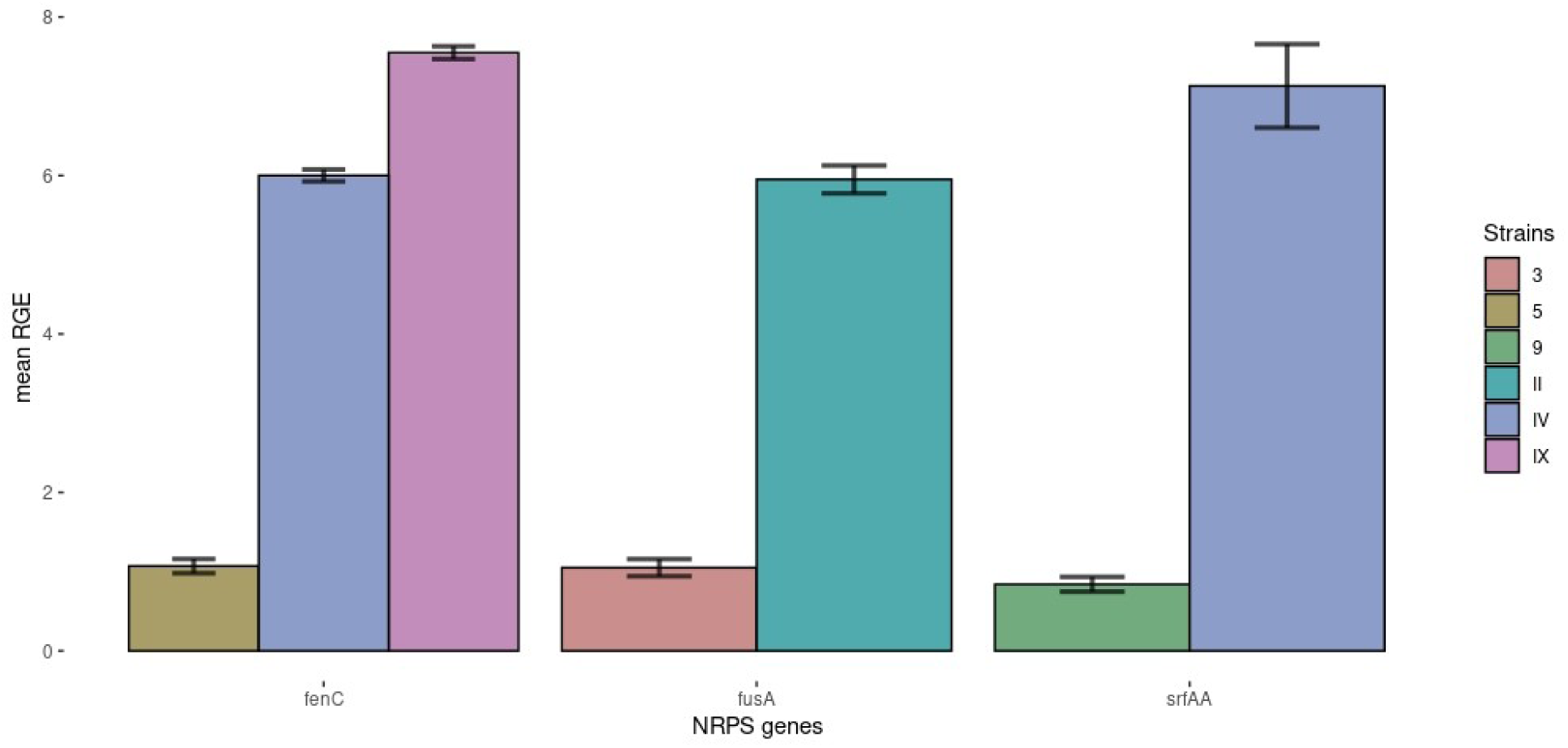
Analysis of relative expression (± SE) of NRPS genes in different bacterial strains in the presence of the fungus *F. oxysporum* Ras6.2.1. Numerical designations on the X-axis: Arabic numerals denote bacterial strains without expressed antagonistic properties but in which NRPS genes were detected. Roman numerals - antagonist bacterial strains.

Subsequently, we assessed the impact of different species of phytopathogenic fungi on alterations in the expression profiles of NRPS genes in selected bacterial strains with robust antagonistic properties.

Fusaricidin synthase was detected in 10 strains, with the expression level of *fusA* varying significantly depending on the co-cultured *Fusarium* strain. Two strains exhibited an increase in *fusA* expression in the presence of fusarium, irrespective of the strain. Specifically, *P. polymyxa* R5.31 demonstrated a 4- and 2.5-fold increase in expression when co-cultivated with Fus 5 and Ras 6.1 (p = 0.0001), while and *B. velezensis* R4.6 displayed a 2- and 12-fold increase in *fusA* expression in a presence of *F. graminearum* Fus5 and *F. oxysporum* Ras6.1. (p = 0.00013). *P. polymyxa* strains R6.14 and K1.14 exhibited increase in expression of *fusA* when co-cultured with Fus5 and a decrease in the presence of Ras6.1(For R6.14 p = 0.005, for K1.14 (p = 0.00064). Conversely, strains R4.5 (p = 0.0001), R4.24 (p = 0.007), O1.27 (p = 0.00016) and V3.14 (p = 0.00081) decreased the expression level of fusaricidin synthase in the presence of Fus5 and increased in the presence of Ras6.1.

Fengycin synthase and surfactin synthase genes were found in *B. velezensis* V3.14 and *B. velezensis* R4.6. The expression level of *fenC* in V3.14 increased almost threefold (p = 0.0001) when co-cultured with Fus5 and decreased twofold (p = 0.0001) when co-cultured with Ras6.1. For strain R4.6, *fenC* expression decreased almost threefold (p = 0.00025) in the presence of Fus5, while co-cultivation with Ras6.1 does not affect the expression level. The changes in surfactin synthase expression were not statistically significant (p = 0.11 for V3.14, p = 0.27 for R4.6).

Polymyxin synthase genes were found in all 8 strains. Among them, only R3.13 had an increased (p = 0.02) expression level in both co-cultivation experiments. Strain R4.24 (p = 0.002) exhibited a fivefold decrease in polymyxin synthase expression level in the presence of Fus5 and a twofold increase in the presence of Ras6.1. Other strains either decrease or show no significant change in the expression level of polymyxin synthase when co-cultured with fusarium strains. Results categorizing the gene expression changes in nonribosomal synthesis genes are visualized in Figures 2-5.

**Fig. 2.**
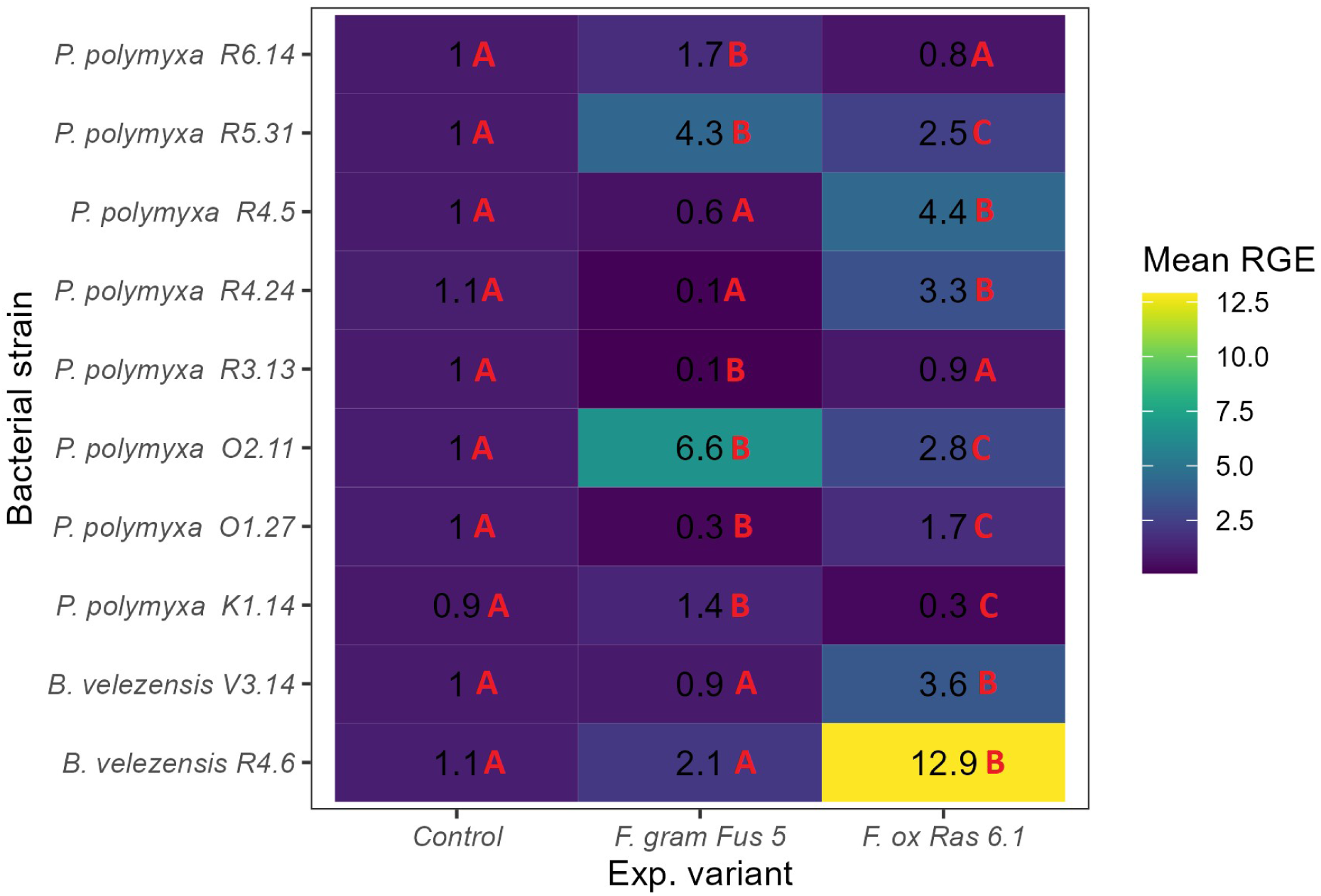
Analysis of relative expression of *fusA* gene in different bacterial strains in the presence of *F. oxysporum* Ras6.1 and *F. graminearum* Fus5. For strain R6.14 F = 15.09, p = 0.005, η^2^G = 0.83; For strain R5.31 F = 140.59, p = 0.0001, η^2^G = 0.98; For strain R4.5 F = 418.06, p = 0.0001, η^2^G = 0.99; For strain R4.24 F = 13.03, p = 0.007, η^2^G = 0.81; For strain R3.13 F = 19.12, p = 0.002, η^2^G = 0.86; For strain O2.11 F = 52.74, p = 0.00016, η^2^G = 0.95; For strain O1.27 F = 46.72, p = 0.00022, η^2^G = 0.94; For strain K1.14 F = 31.76, p = 0.00064, η^2^G = 0.91; For strain V3.14 F = 29.14, p = 0.00081, η^2^G = 0.91; For strain R4.6 F = 56.51, p = 0.00013, η^2^G = 0.95.

**Fig. 3.**
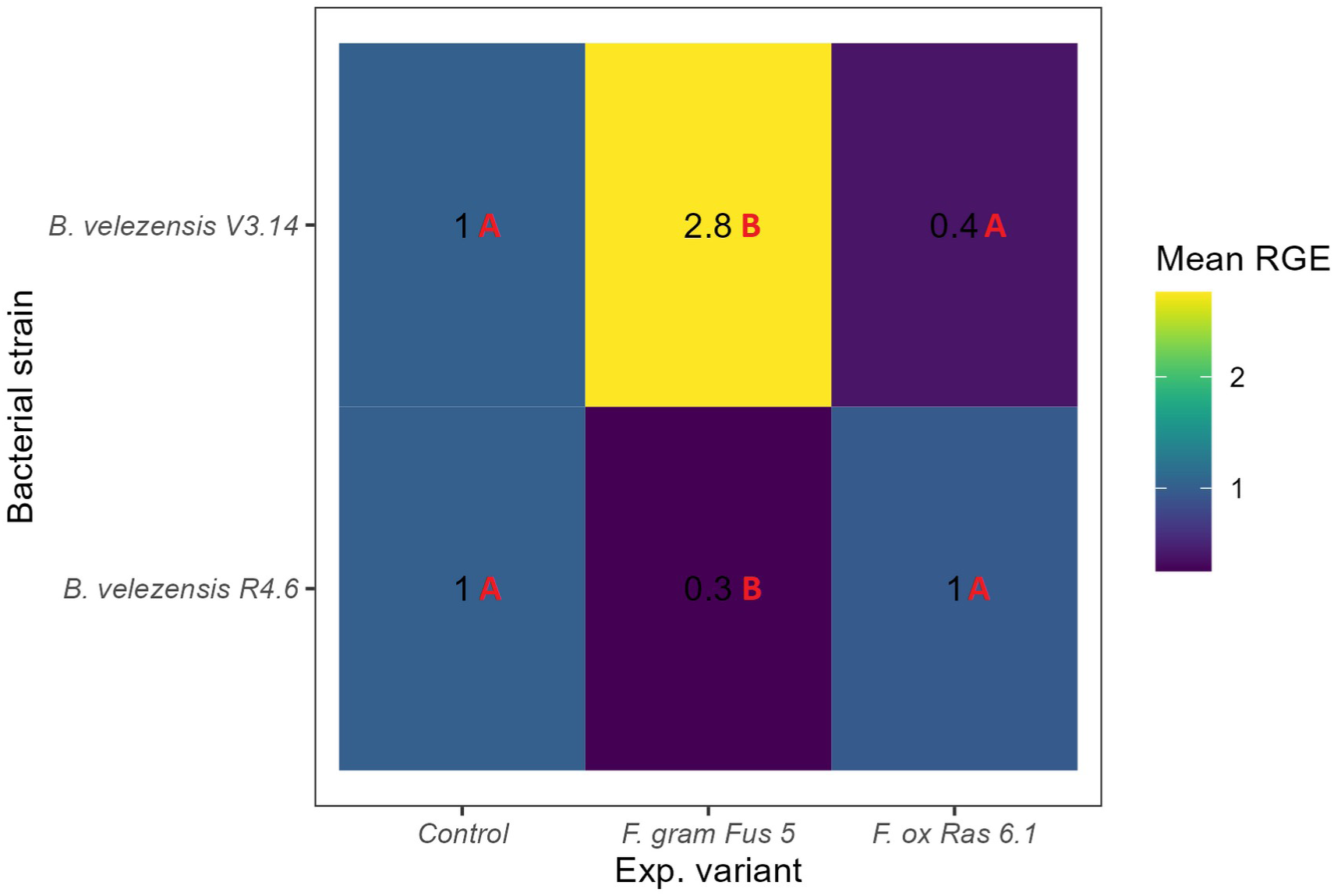
Analysis of relative expression of *fenC* gene in different bacterial strains in the presence of *F. oxysporum* Ras6.1 and *F. graminearum* Fus5. For strain V3.14 F = 63.79, p = 0.0001, η^2^G = 0.96; For strain R4.6 F = 44.81, p = 0.00025, η^2^G = 0.94.

**Fig. 4.**
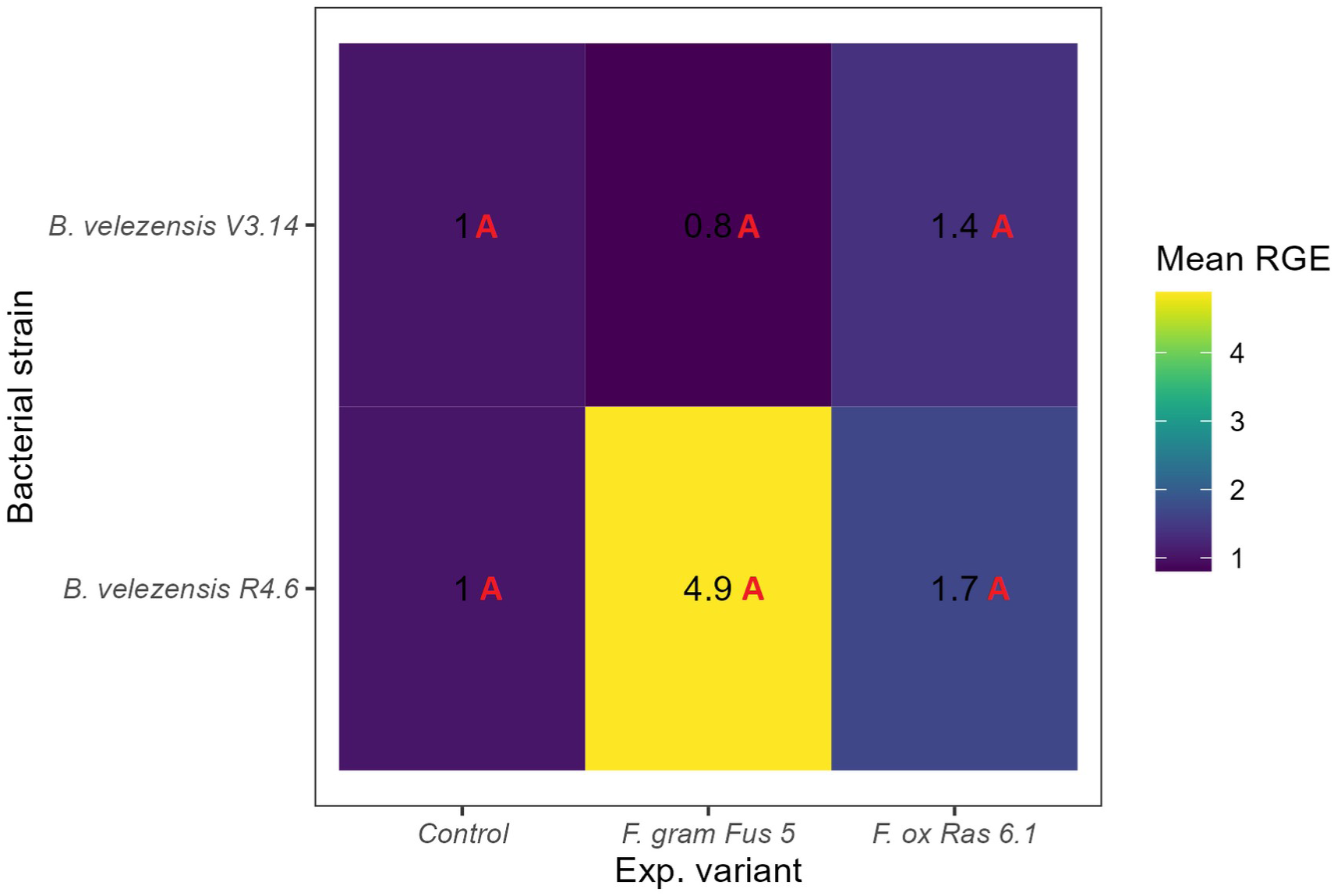
Analysis of relative expression of *srfAA* gene in different bacterial strains in the presence of *F. oxysporum* Ras6.1 and *F. graminearum* Fus5. For strain V3.14 F = 3.35, p = 0.11, η^2^G = 0.53; For strain R4.6 F = 1.64, p = 0.27, η^2^G = 0.35.

**Fig. 5.**
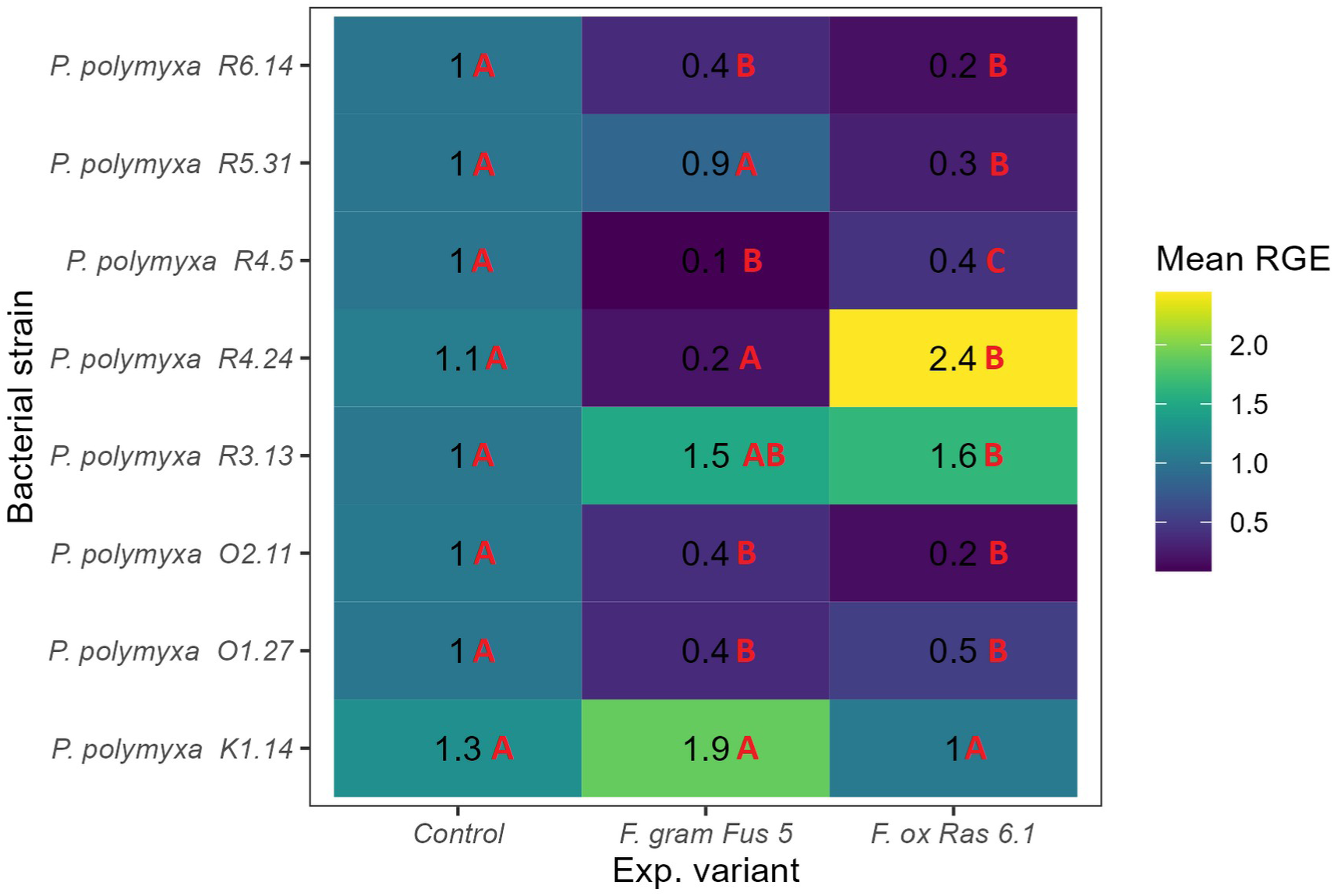
Analysis of relative expression of *pmxE* gene in different bacterial strains in the presence of of *F. oxysporum* Ras6.1 and *F. graminearum* Fus5. For strain R6.14 F = 81.57, p = 0.0001, η^2^G = 0.96; For strain R5.31 F = 72.6, p = 0.0001, η^2^G = 0.96. For strain R4.5 F = 73.43, p = 0.0001, η^2^G = 0.96; For strain R4.24 F = 19.78, p = 0.002, η^2^G = 0.87. For strain R3.13 F = 8.13, p = 0.02, η^2^G = 0.73; For strain O2.11 F = 10.86, p = 0.01, η^2^G = 0.78. For strain O1.27 F = 21.07, p = <0.002, η^2^G = 0.88; For strain K1.14 F = 5.04, p = 0.052, η^2^G = 0.63.

### 2.3. Field trials of the bacterial preparation

Field trials of the bacterial preparation were conducted to assess its impact on *Fusarium* fungi in sugar beet crops and soil. Results from field 1 showed no significant difference in *Fusarium* abundance in root crops after a single treatment compared to the control group (p=0.54). However, with a double treatment, there was a statistically significant 44% decrease (p=0.048). In the soil under sugar beet on the same field, single treatment did not lead to significant differences in *Fusarium* numbers (p=0.18), while double treatment led to a significant 3.8-fold reduction (p=0.005).

Field 2 displayed widespread *Fusarium* presence in sugar beet crops in the control group. Both single and double treatments with the preparation statistically significantly decreased the fungi population (p=0.005 and p=0.023, respectively). Similar trend was observed in the soil under the beet at this site as in root crops, where only double treatment significantly reduced the fungi abundance compared to the control group (p=0.048).

In field 3, *Fusarium* levels in sugar beet root crops ranged from medium to low, and both single and double treatments resulted in a significant 3-4 times reduction in the fungus population (p=0.004 and p=0.01, respectively). In the soil at this site, there was a 4.5-6.7-fold decrease in *Fusarium* abundance compared to control. In single and double treatment experiments, p-values were 0.0013 and 0.003, respectively.

Field 4 demonstrated significant reductions in *Fusarium* occurrence of in root crops, more than halved, only after double treatment (p=0.035). Single treatment did not show significant differences (p=0.17). In the soil at this site, both single and double treatments significantly reduced *Fusarium* abundance, more than halved, compared to control group (p=0.006 and p=0.001, respectively). These results are summarized in Figure 6.

**Fig. 6.**
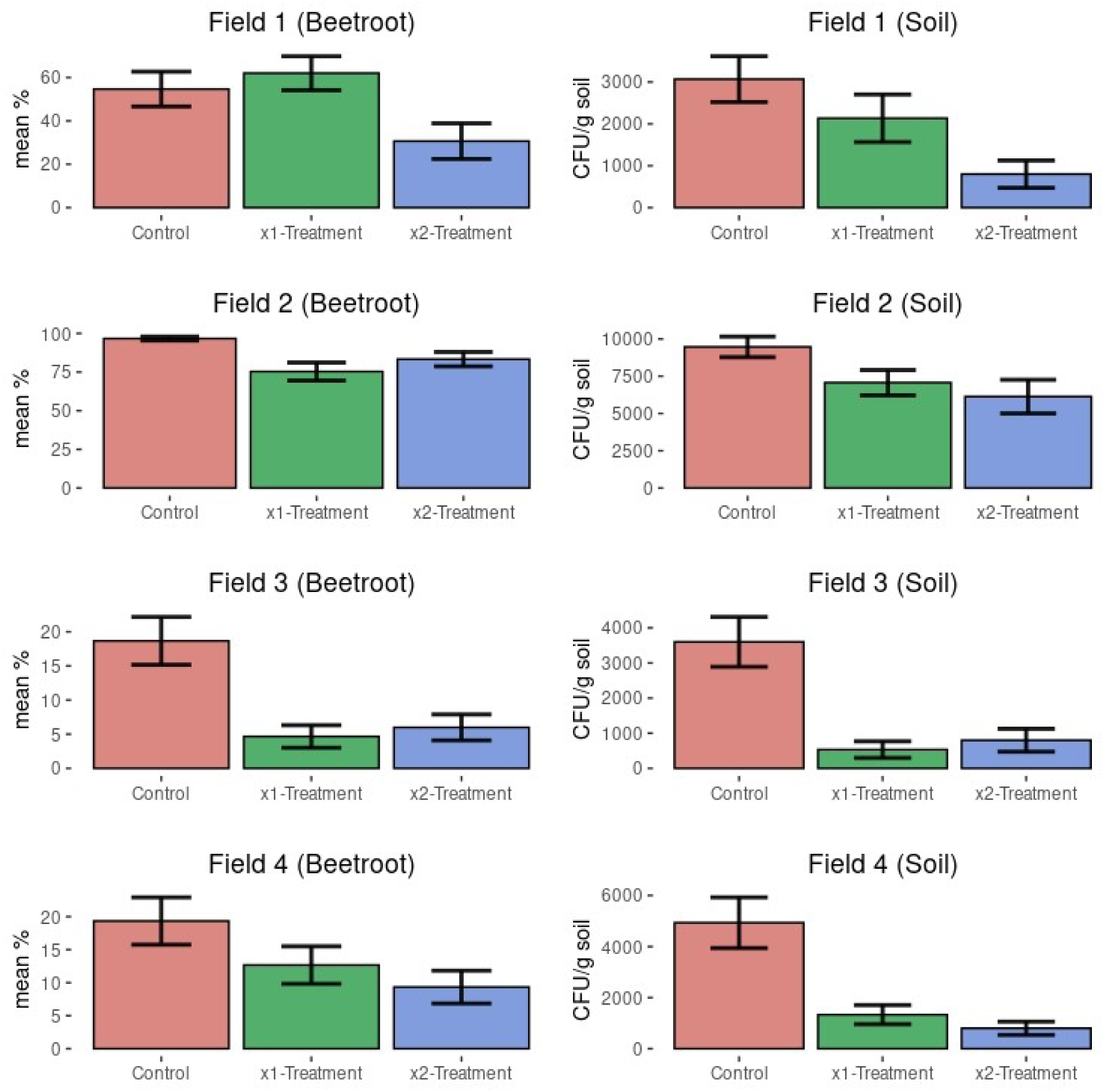
Summarized results of field trials of biopreparation based on 10 studied bacterial strains. Antifungal activity is expressed in % of beetroot fragments that give fusarium growth when inoculated on dense nutrient medium, as well as by its abundance in the soil, which is expressed in the number of CFU/g of soil. Data are given in the form of mean value and standard error.

## 3. Discussion

Our study revealed the absence of chitinase activity in all strains studied, which implies that their antifungal activity is due to the work of alternative mechanisms. These 10 new fusarium antagonist bacterial strains antagonistic activity is attributed, apparently, mainly to nonribosomal synthesized metabolites, as confirmed by the PCR-analysis of target genes involved in nonribosomal synthesis. In turn, only 3 strains with weak antifungal activity (strains 3,5,9) were found to have NRPS genes, with their NRPS expression levels significantly lower than those of highly antagonistic strains. However, it is worth noting that besides chitinase and NPR, bacillus are capable of producing some other antifungal compounds related to volatile organic compounds and siderophores that were not considered in our study (Bharucha Ud et al., 2013; Ghazy and El-Nahrawy, 2021; Saravanan R et al., 2021).

The 10 antagonist bacterial strains used in this study were identified as strains of *Paenibacillus polymyxa* (strains K1.14, O1.27, O2.11, R3.13, R4.5, R4.24, R5.31, R6.14) and *Bacillus velezensis* (V3.14, R4.6). The spectrum of antifungal NRPs that can be produced by these strains and against 2 different *Fusarium* species was analysed. The highest number of NRPS genes was found in *B. velezensis* V3.14 and R4.6 strains. Upon analysis of PCR products with specific primers, the presence of surfactin and fengycin synthetases genes, along with closely related fusaricidin synthetase genes, was detected in these strains. Furthermore, genes responsible for fusaricidin and polymyxin synthetase synthesis were detected in all *P. polymyxa* strains.

The expression level of the gene responsible for fusaricidin synthesis in the vast majority of bacterial antagonist strains were found to be elevated when co-cultured with the Ras6.1. Conversely, only one bacterial strain out of 10 (V3.14) exhibited increased expression of fengycin, and only in the presence of fungus Fus5. This suggests that these antagonist bacterial strains can produce the specific agent against a particular pathogen in its presence. For instance, fusaricidin is predominantly produced in response to *F. oxysporum*, while in the case of *F. graminearum*, it is more often fengycin. Cases where both fusaricidin and fengycin expression decreased in the presence of either fungi may indicate that these strains are capable of synthesizing other nonribosomal peptides or antagonistic agents, and their genome needs to be checked for new NRPS genes.

Assessing the number of expressed NRPS genes in the bacterial strains presented in this work, we noted a variation across strains. Some strains, such as O1.27, R4.5, R6.14, expressed a single NRPS gene responsible for fusaricidin synthetase, while others (R5.31, O2.11, R3.13, R4.24) expressed 2 genes - from polymyxin and fusaricidin synthetase clusters. Two strains, V3.14 and R4.6, exhibited altered expression of genes responsible for fusaricidin and fengycin synthesis. Notably, V3.14 showed induction of fusaricidin gene expression and suppression of fengycin gene expression in response to the presence of the Ras6.1 fungus, but the opposite pattern emerged when co-cultured with the fungus Fus5. In contrast, strain R4.6 displayed increased fusaricidin gene expression when co-cultured with both types of fungi.

It seems that each individual strain responded differently to the presence of a particular species of *Fusarium*, which may indirectly indicate the use of different strategies by the bacteria against different fungal species.

The differential gene expression of nonribosomal peptide synthetase genes differed several times compared to control levels. The discrepancies observed between strains may be arise from both differences in the NRPS gene clusters themselves and differences in the function and structure of regulatory genes (e.g. *degQ*, *abrB*, etc.) (Lilge et al., 2021; Vasilchenko et al., 2022).

Moreover, these differences between strains could be influenced by epigenetic regulation of gene expression as recent discoveries have highlighted distinct mechanisms of epigenetic regulation impacting gene expression within this group (Vasilchenko et al., 2022).

Notably, strain R3.13 showed increased polymyxin expression in response to Fus5 and Ras6.1 presence (1.5- and 1.6-fold, respectively), with a concurrent decrease in fusaricidin expression (0.9- and 0.1-fold, respectively). While polymyxin is typically recognized as an inhibitor of bacterial growth, its elevated expression in this scenario may suggest a role in suppressing fungal growth. It is also plausible that the NRPS responsible for polymyxin synthesis participates in the assembly of another compound active against fungi, indirectly elucidating the decrease in fusaricidin expression.

In the light of the identified strain specificity, a reasonable strategy was to develop a complex preparation composed of different strains, assuming that they would complement each other’s activity. Consequently, the biopreparation based on 10 studied bacterial strains was obtained.

Evaluating the biopreparation’s efficacy across various farms revealed significant variations. For instance, for field 1, significant differences were observed in both root crops and soil when treated twice, and for field 2, both single and double treatments significantly reduced the fungal population in root crops, while in soil only double treatment significantly reduced the fungal population. Fields 3 and 4 there exhibited significant reductions in fusarium abundance in soil both in double and single treatments, while in field 4 a statistically significant reduction of infection background in root crops was observed only in double treatment. In field 3 the infectious background of fusarium in root crops was significantly reduced in both single and double treatments. Therefore, double treatment consistently demonstrated more pronounced differences from the control in decreasing fungal populations compared to single treatments.

It seems that the ability to increase the production of specific fungicides in response to the particular fungi may be a promising marker to select the most effective strains. Moreover, species- and strain-specificity of bacterial response to different antagonist fungi is a crucial factor to consider in the development and utilization of antifungal biopreparation.

In the realm of preparations development, at least two approaches are to be explored: firstly, the design of bioagents tailored precisely to specific phytopathogens, and secondly, the formulation of a mix of highly targeted strains, that will be effective against multiple pathogens simultaneously. In this study, we utilized phytopathogenic fungi sourced from the same region as our research site, elevating hopes for the drug’s efficacy. Nevertheless, in agricultural practice, the potential emergence of novel strains, mutations, horizontal gene transfers, and the ingress of additional phytopathogen species and strains from neighbouring fields remain as possible challenges. Moreover, the swift and accurate identification of a particular pathogen may not always be feasible. Hence, we believe that the most promising strategy involves the development of a composite mixture of highly specific, inducible by pathogen bacteria.

Further research avenues could explore the range of metabolites and toxins released by diverse *Fusarium* strains, shedding light on the stimuli influencing bacterial strain adaptation against specific pathogens.

